# Serological evidence of exposure to Rift Valley, Dengue and Chikungunya Viruses among agropastoral communities in Manyara and Morogoro regions in Tanzania: A community Survey

**DOI:** 10.1101/2020.01.16.908830

**Authors:** Rule M. Budodo, Pius G. Horumpende, Sixbert I. Mkumbaye, Blandina T. Mmbaga, Richard S. Mwakapuja, Jaffu O. Chilongola

**Author notes:** Correspondences (JC). These authors contributed equally to this work (Rule M. Budodo, Pius G. Horumpende).

## Abstract

**Introduction:** Tanzania has recently experienced outbreaks of dengue in two coastal regions of Dar es Salaam and Tanga. Chikungunya and Rift Valley Fever outbreaks have also been recorded in the past decade. Little is known on the burden of the arboviral disease causing viruses (Dengue, Rift Valley and Chikungunya) endemically in the inter-epidemic periods. We aimed at determining the prevalence of the dengue, rift valley and chikungunya among humans in two geo ecologically distinct sites.

**Methodology:** The community-based cross-sectional study was conducted in Magugu in Manyara region and Mvomero in Morogoro region in Tanzania. Venous blood was collected from participants of all age groups, serum prepared from samples and subjected to ELISA tests for RVFV IgG/IgM, DENV IgG/IgM, and CHIKV IgM/IgG. Samples that were positive for IgM ELISA tests were subjected to a quantitative RT PCR for each virus. A structured questionnaire was used to collect socio-demographic information. Data analysis was conducted using SPSSv22.

**Results:** A total of 191 individuals from both sites participated in the study. Only one CHIKV was detected in Magugu site but none of the 69 participants from Magugu site was seropositive or positive for RVFV and DENV. Of the 122 individuals from Wami-Dakawa site, 16.39% (n=20) had recent exposure to RVFV while 9.83% (n=12) were recently infected by Chikungunya virus. All samples were negative by RVFV and CHIKV qPCR. Neither Infection nor exposure to DENV was observed in participants from Wami-Dakawa. Risk factors associated with RVFV and DCHIKV seropositivity were being more than 5 in a household, having no formal education and having recently travelled to an urban area.

**Conclusion:** We report an active circulation of RVFV and CHIKV in humans in Wami-Dakawa, in Mvomero district of Morogoro region during dry season, a higher rate of exposure to RVFV compared to CHIKV and an absence of circulating RVFV, DENV and CHIKV in humans in Magugu site in Manyara region.

**Author Summary:** Dengue, Chikungunya, rift valley and other viruses constitute an important group of etiologies of fever in Tanzania. In the past decade, Tanzania has experienced DENV and RVFV at different times. While RVFV outbreaks have occurred cyclically in approximately ten-year periods in Tanzania, DENV outbreaks have been more frequent since 2010. CHIKV infection is an important but largely unrecognized illness in Tanzania. In this study, we aimed to generate baseline data on the exposure and infection status of DENV, RVFV and CHIKV by detecting antibodies to the viruses and detecting the viruses in human subjects in two geo-ecological distinct sites. Neither infection nor exposure to the viruses were in observed in Magugu site in Manyara region, northern Tanzania. There was a significant exposure to RVFV (16.39%) and CHIKV (9.83%) in Wami-Dakawa but not to DENV in either site. None of the viruses was detected by PCR in any of the sites. Potential risks for exposure to CHIKV and RVFV were Larger numbers of household members, having no formal education and having recently travelled to an urban destination. Since arbovirus outbreaks are usually unpredictable, it is crucial to undertake active surveillance for RVFV, DENV, CHIKV and other viral agents in Tanzania.

## Introduction

Rift valley fever virus (RVFV), Dengue virus (DENV) and Chikungunya virus (CHIKV) are endemic in Sub-Saharan Africa and cause sporadic and sometimes large epidemics in humans. The ecological drivers of the pattern and frequency of virus infections (and subsequent epidemics) in the different host species are largely unknown [1]. Many authors identified non-human primates, mammalian animals and birds as hosts/reservoirs for arboviruses [2-4]. Although Arboviruses are genomically variant, they share a common transmission mode through vectors, pathobiological mechanisms and cause overlapping clinical presentations [5]. DENV and CHIKV are increasing global public health concerns due to their rapid geographical spread and increasing disease burden.

The transmission of these viruses is complicated by human population migration/resettlement, internal displacement due to environmental and civil unrest in many African countries and the triad of the modern world: urbanization, globalization, and international mobility (Kraemer et al., 2015). CHIKV causes large epidemics with serious economic and social impact and its clinical presentations are similar to several flavivirus infections [6-9]. Dengue virus has caused major epidemics in past centuries. In the Americas, dengue was effectively controlled in the mid-1900s by effective control of *A. aegypti*, the principal urban vector of both viruses. However, there has been a re-emergence of dengue worldwide [10]. Previous studies demonstrated that some degree of DENV genetic variability is necessary for the persistence during periods of endemicity and epidemic outbreaks [11]. For decades, arboviral diseases were considered to be of minor concern with regard to global mortality and disability. Consequent to this, low priority has been given to arbovirus research and related public health infrastructure. In the past decade, however, an unprecedented emergence of epidemic arboviral diseases (notably dengue, chikungunya, and Rift valley fever) has been recorded in Tanzania. It is important to understand the prevalence of arboviruses in different weather conditions in Tanzania in order to ascertain their presence in different hosts at different weather conditions annually.

Arboviruses are primarily maintained by horizontal transmission (HT) between arthropod vectors and vertebrate hosts in nature. Occasionally, they transmitted vertically in the vector population from an infected female to the offspring, which is a proposed maintenance mechanism during adverse conditions [7]. For RVFV, it is known that factors such as dense vegetation, suitable temperature conditions, and the presence of ruminants make it favorable for mosquitoes to breed, replicate the virus, and pass it on to animals and humans. The recent epidemics caused by these arboviruses have been associated with many factors including urbanization/population growth and international travel and trade, allowing for spread of vectors, and spreading of arboviruses into new niches, followed by amplification through human-vector-human cycle. Climate change is predicted to further impact on the distribution of vector-borne diseases, such as Rift Valley fever and dengue, which are highly sensitive to climatic conditions [12]. The maintenance mechanisms during inter epidemic periods (IEPS) become interesting as to where the viruses hide during the “silent” periods. This study was therefore carried out to identify the burden of arboviruses in two geo-ecologically different sites in North Eastern and Central Eastern Tanzania during the dry season to generate baseline data that will form the basis of a future ‘across-season’ transmission study.

## Material and Method

### Study design, Study sites and sample size estimation

This was a cross sectional, community-based survey conducted in two sites, namely; Magugu Ward of Babati Rural district in Manyara region and Wami-Dakawa ward in Mvomero district in Morogoro region. The study involved a total of 191 participants in both study sites, 69 participants from the Magugu site whereas 122 participants were from Wami-Dakawa site.

Magugu site (S3099’ S4001’; E35070’ E35077’) is located in Babati Rural District, in northern Tanzania along the rift valley, at approximately 900m to 1200m above sea level. Magugu Ward has seven villages and a total population of 26,131 people. The main economic activities include crop agriculture and livestock keeping. There is one government owned health center which serves all the seven villages. Average annual rainfall is about 650mm. There are two rainy seasons, short rainy season from October to December and long rainy season from mid-March to May, followed by a cool and dry season from June to mid-August, and a hot dry period from mid-August to October.

Wami-Dakawa is a ward in Mvomero district of Morogoro region. The site is It is bordered to the north by the Tanga Region, to the northeast by the Pwani (Coast) Region, to the east and southeast by Morogoro Rural District and Morogoro Urban District and to the west by Kilosa District. According to the 2012 Tanzania National Census, the population of Mvomero District is estimated at312,109 [13]. The district lies between latitudes 6° south and 8° south and longitudes 36° 30′ east and 38° east. The area has semi humid climate with an average annual rainfall of 800 mm. The short rains start in November and end in January followed by heavy rainfall between March and May. The district experiences a dry season from June to October and the average annual temperature is 24.6°C. The district has an area of 14,245 square kilometers and a population of 438,175 people [14]. In addition, outbreaks of Rift Valley fever (RVF) have been reported in the district and the most recent outbreak in 2007 was associated with human and livestock morbidity as well as significant mortalities and abortions in livestock. However, there are no reports describing the occurrence of arbovirus infections, such as DENV, CHKV, and RVFV [9].

### Blood ample collection, processing and storage of specimens

Venous blood samples were collected from median cubital vein following venipuncture. One and a half ml of the blood sample was placed in plain clot activating tubes for serum preparation. The remaining 1.5 ml was placed into EDTA tubes and an equal volume of Trizol added as per manufacturer’s instructions (Zymo Research, Irvine, CA, U.S.A.). The mixture was gently mixed for up to 5 minutes and immediately both types of samples were kept at 4°C in cool boxes before they were shipped to the laboratory. A standardized questionnaire was used to collect clinical and socio-demographic information from participants.

### Laboratory procedures

#### DENV and CHIKV IgG and IgM ELISAs

Briefly, serum from plain tubes was obtained by centrifugation of samples at 2,000 rpm x g for 10 minutes in a refrigerated centrifuge. Serum samples were stored at −20°C awaiting serological analyses. For seropositivity of CHIKV, anti-CHIKV IgM and IgG were analyzed using an Indirect ELISA kit (SD, Gyeonggi-do, Korea and IBL international, Hamburg, Germany, respectively). Detection of DENV IgG and IgM antibodies were done using a direct enzyme linked immunosorbent assay (ELISA) kit (SD Inc, Gyeonggi-do, Korea) as described by [15]. All assays were performed according to manufacturers’ instructions. Optical density (OD) reading was done at 450 nm and the units of antibody concentration and cut-off values calculated as described by the manufacturers. Briefly, for the Anti-DENV IgM/IgG and IgM anti-CHIKV ELISAs the diagnostic cut-off value was calculated as the average OD of negative controls + 0.300. For the IgG CHIKV ELISA the threshold for positivity was based on the OD cut-off value of the cut-off control + 10 %.

#### RVFV competitive ELISA

All samples were selected for testing for antibodies against RVFV using a competitive ELISA (ID Screen Rift Valley Fever Competition Multi Species, ID-vet, Grables, France), which detects both IgG and IgM antibodies directed against the nucleoprotein of RVFV. Validation tests for the test has shown to have a sensitivity of between 91 and 100% and a high specificity (100%). In both tests done by the manufacturer and an independent ring trial [16]. The competitive ELISA was performed according to the instructions of the manufacturer and as described previously [17]. In order to control the validity of each plate, the mean value of the two negative controls (OD_NC_) was computed whereby a plate was considered valid if the OD_NC_ was>0.7. For a valid plate, the mean value of the two positive controls divided by OD_NC_ had to be <0.3. For each sample, the competition percentage was calculated by dividing OD_sample_/OD_NC_) × 100. If the value was equal or less than 0.4, the sample was considered positive while a value greater than 50% was considered negative.

#### Ribonucleic acid extraction and RT-PCR procedures

For DENV and CHIKV, Blood samples kept in EDTA tubed were centrifuging at 1,000 rpm x g for 10 minutes in a refrigerated centrifuge to obtain buffy coat. Ribonucleic acid (RNA) was extracted from buffy coat samples using the Boom method [18]. cDNA were synthesized using Superscript^®^ VILO™ cDNA synthesis kit (Invitrogen, life technologies, USA) according to manufacturer’s instructions and PCR done as previously described [15]. For RVFV detection, RNA was extracted from sera using a QIAamp viral RNA mini kit (QIAGEN, Germany) as per manufacturer’s instructions. RVFV RNA was detected using TaqMan probe-based one-step RT-PCR targeting the RVFV Gn gene as described by Gudo and colleagues [18].

#### Nature of data and data Analysis

All data were collected strictly following GCP guidelines. Participants were identified by anonymous IDs and initials as the only way of their identification. The main tool for data collection was the case report form (CRF). Data that were collected through the CRFs were demographics, characteristics of households, characteristics of livestock herds own by participants, treatment history, travel, co-morbidities, virologic and serological parameters. Data were analyzed using SPSS v.22 (IBM^®^ Corp., Armonk, NY, USA). Descriptive data are reported as frequencies, means and medians. Categorical data are reported as tabulation of proportions. Logistic regression analyses were used to examine associations between seropositivity to RVFV, DENV and CHIKV and candidate predictor variables. Bivariate models were constructed for candidate demographic variables and herd characteristics evaluated. All variables with a coefficient p-value ≤0.2 in the bivariate model were considered in the multivariate modelling. Both crude odds ratios (cORs) from bivariate models and adjusted odds ratios (aORs) from the multivariate model are reported.

#### Ethical considerations

Ethical clearance was sought from Kilimanjaro Christian Medical University College Research and Ethics Review Committee (CRERC) and clearance certificate number 2419 was obtained. Permission to conduct the study was sought from both Manyara and Morogoro Regional Administrative Secretaries. The Babati Rural and Mvomero District Executive Officers and District Medical Officer were consulted for permission.

## Results

### Descriptive statistics

A total of 191 sample were collected, 122 from Mvomero and 69 from Magugu. None of the viruses was detected either by serology or by PCR among the 69 samples collected from Magugu site. Due to the clear absence of any positive test results in Magugu, presented results include analyses for samples collected from Wami-Dakawa, in Mvomero district. Of the 122 individuals participated in the study from Wami-Dakawa site, 58 (47.5%) individuals were involved in pastoralism as their main occupation while 64 (52.5%) were peasant farmers. About two thirds of the participants 80 (65.6%) live in households with less than 5 family members. About three-fifth (60.7%) of the interviewed participants, reported not to have travelled outside Wami-Dakawa during the previous three months, with 33/48 (68.8%) of those who had travelled having travelled to an urban destination. A total of 60 (49.2%) participants had herd sizes made of less or equal 30 goats. Goat browsing was mainly done in communal grasslands by 64.8 % of the participants. Browsing was done about 70 km from Mikumi national park/game reserves, with minimal interaction between livestock and wild animals. (Table 2). Out of the 122 people tested for RVFV, 20 (16.39%) were positive for IgM/IgG antibodies to RVFV, this indicate that there is active circulation of RVFV in the area. Twelve of the participants (9.83%); tested positive for anti-CHIKV IgM/IgG (Table3). Only one participant was seropositive for CHIKV IgG from Magugu site. Through interviews, the patient who was seropositive for CHIKV IgM/IgG reported to have arrived in Magugu 3 weeks ago from Tanga region. All samples that were RVFV and CHIKV IgM/IgG positive were negative for qPCR.

**Table 1:**
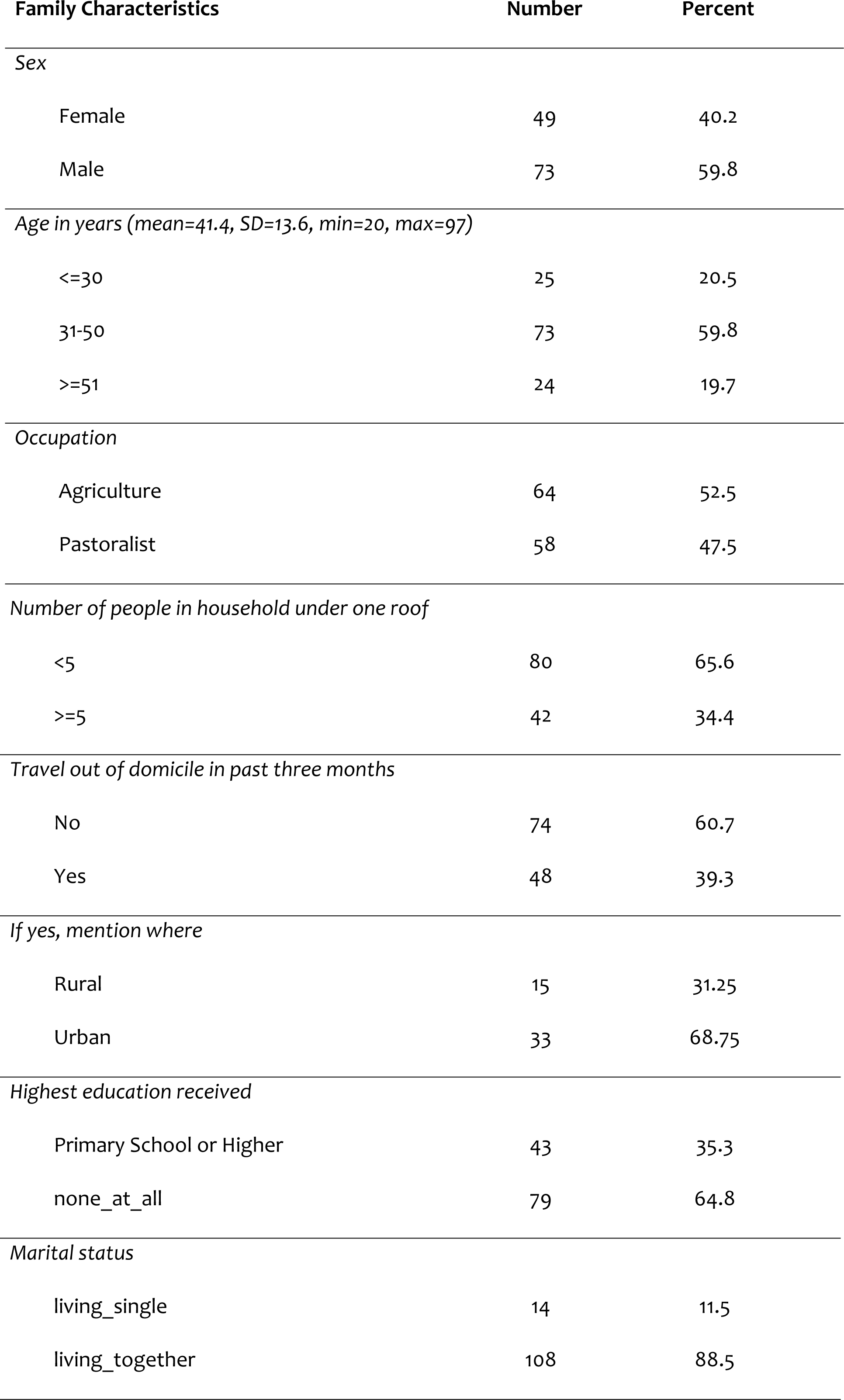
Demographic characteristics of participants.

| Family Characteristics | Number | Percent |
| --- | --- | --- |
| *Sex* |  |  |
| Female | 49 | 40.2 |
| Male | 73 | 59.8 |
| *Age in years (mean=41.4, SD=13.6, min=20, max=97)* |  |  |
| <=30 | 25 | 20.5 |
| 31-50 | 73 | 59.8 |
| >=51 | 24 | 19.7 |
| *Occupation* |  |  |
| Agriculture | 64 | 52.5 |
| Pastoralist | 58 | 47.5 |
| *Number of people in household under one roof* |  |  |
| <5 | 80 | 65.6 |
| >=5 | 42 | 34.4 |
| *Travel out of domicile in past three months* |  |  |
| No | 74 | 60.7 |
| Yes | 48 | 39.3 |
| *If yes, mention where* |  |  |
| Rural | 15 | 31.25 |
| Urban | 33 | 68.75 |
| *Highest education received* |  |  |
| Primary School or Higher | 43 | 35.3 |
| none_at_all | 79 | 64.8 |
| *Marital status* |  |  |
| living_single | 14 | 11.5 |
| living_together | 108 | 88.5 |

**Table 2:** Characteristics and grazing practices of participants.

| **Herd Characteristics** | **Number** | **Percent** |
| --- | --- | --- |
| *Herd size* |  |  |
| <=30 | 60 | 49.2 |
| 31-50 | 12 | 9.8 |
| >50 | 50 | 41.0 |
| *Characteristic of grazing area* |  |  |
| Forest | 41 | 33.6 |
| Plain grasslands | 79 | 64.8 |
| Savannah | 2 | 1.6 |
| *Do you graze very close to game reserve* |  |  |
| No | 121 | 99.2 |
| Yes | 1 | 0.8 |
| *Do you camp your animals during dry season* |  |  |
| No | 104 | 85.3 |
| Yes | 18 | 14.8 |
| *Do you meet wildlife where you take your herd for water drinking?* |  |  |
| No | 103 | 84.4 |
| Yes | 19 | 15.6 |
| *Distance of grazing in kilometers* |  |  |
| <=10 | 92 | 75.4 |
| Nov-20 | 4 | 3.3 |
| >=31 | 26 | 21.3 |
| *Distance of water source in kilometers* |  |  |
| <=10 | 94 | 77.1 |
| Nov-20 | 2 | 1.6 |
| >=31 | 26 | 21.3 |
| *Distance from wildlife in kilometers* |  |  |
| <=10 | 61 | 50.0 |
| Nov-20 | 1 | 0.8 |
| 21-30 | 22 | 18.0 |
| >=31 | 38 | 31.2 |

**Table 3:** Enzyme Linked Immunosorbent Assay results for IgM/IgG antibodies to Rift Valley Fever Virus and Chikungunya Viruses.

| Total<br>tested | Rift Valley Fever IgG/IgM seropositivity |  |  |  | Chikungunya IgM/IgG seropositivity |  |  |  |  |
| --- | --- | --- | --- | --- | --- | --- | --- | --- | --- |
|  | +ve | % | 95% C.I. |  | Total<br>tested | +ve | % | 95% C.I. |  |
|  |  |  | Low | Up |  |  |  | Low | Up |
| 122 | 20 | 16.39 | 9.8 | 23.6 | 122 | 12 | 9.83 | 4.9 | 15.7 |

All Samples were qPCR negative for RVFV, DENV and CHIKV; only one CHIKV case was detected by ELISA from Magugu site in a participant just travelled from Tanga.

### Risk factors for Rift Valley Fever Virus IgG/IgM seropositivity

Participants aged between 31 and 50 were less likely to be seropositive for RVFV IgG/IgM [OR 0.64; (95%CI: 1.8-2.33), p<0.05]. Households with more than 5 people sleeping in the same house were 2.15 times more likely to be RVFV IgG/IgM seropositive compared to households with less than 5 people [OR 2.15, (95%CI: 2.43-5.1), p<0.05]. Similarly, participants without formal education had higher odds of being RVFV IgG/IgM seropositive compared to participants with formal education [OR 1.9, (95%CI: 2.63-4.56), p<0.05]. (Table 4).

**Table 4:** Risk Factors for RVFV IgG/IgM Seropositivity.

| Predictors of IgG/IgM | OR | P-value | 95% C.I. |  |
| --- | --- | --- | --- | --- |
|  |  |  | Lower | Upper |
| Age in years |  |  |  |  |
| <=30 | 1 |  |  |  |
| 31-50 | 0.64 | 0.050 | 1.8 | 2.33 |
| >=51 | 1.35 | 0.678 | 0.33 | 5.56 |
| Occupation |  |  |  |  |
| agriculture | 1 |  |  |  |
| agropastoralist | 0.77 | 0.608 | 0.29 | 2.08 |
| Place travelled |  |  |  |  |
| Rural | 1 |  |  |  |
| Urban | 1.68 | 0.553 | 0.30 | 9.34 |
| no of people |  |  |  |  |
| <5 | 1 |  |  |  |
| >=5 | 2.15 | 0.038 | 2.43 | 5.10 |
| Education |  |  |  |  |
| Primary school or higher | 1 |  |  |  |
| none_at_all | 1.90 | 0.029 | 2.63 | 4.56 |

### Risk factors for Chikungunya IgM/IgG seropositivity

Results in table 5 show that, participants who had recently travelled to an urban place were 3 times more likely to be seropositive for CHIKV IgG/IgM antibody as compared to participants who had travelled to a rural destination [OR 3.0, (95%CI: 3.3 −27.6), p<0.05]. Household with 5 or more members living in the same house were more likely to be seropositive for CHIKV IgG/IgM antibody as compared to households with less than 5 people [OR 3.5, (95%CI: 1.6-4.84), p<0.05]. Likewise, participants with lack of formal education had higher odds of being seropositive for CHIKV compared to those with formal education [OR 2.85, (95%CI: 2.4 −5.0), p<0.05].

**Table 5:** Risk factors for Chikungunya IgM Seropositivity.

| Predictors of Chikungunya | OR | P-value | 95% C.I. |  |
| --- | --- | --- | --- | --- |
|  |  |  | Lower | Upper |
| Age in years |  |  |  |  |
| <=30 | 1 |  |  |  |
| 31-50 | 1.25 | 0.788 | 0.25 | 6.37 |
| >=51 | 0.83 | 0.861 | 0.11 | 6.46 |
| Occupation |  |  |  |  |
| agriculture | 1 |  |  |  |
| Agro-pastoralist | 1.12 | 0.855 | 0.34 | 3.68 |
| Place travelled |  |  |  |  |
| rural | 1 |  |  |  |
| urban | 3.00 | 0.033 | 3.3 | 27.60 |
| no of people |  |  |  |  |
| <5 | 1 |  |  |  |
| >=5 | 3.5 | 0.033 | 1.6 | 4.84 |
| Education |  |  |  |  |
| Primary school or higher | 1 |  |  |  |
| none_at_all | 2.85 | 0.041 | 2.4 | 5.00 |

## Discussion

The aim of this study was to investigate the prevalence of antibodies against DENV, CHIKV and RVFV in humans and to detect the presence of the viruses in the human hosts. The study also pursued to identify potential predictors for infection by the viruses and seropositivity to the viruses during the inter-epidemic period (IEP). Results indicate that neither infection nor exposure to the viruses among the 69 samples tested in Magugu site in Manyara region. However, the results indicated that significant number of people were infected with RVFV and CHIKV in Wami-Dakawa though it was in dry season with low mosquito challenge. None of the participants were positive for DENV.

Tanzania has experienced outbreaks due to DENV and RVFV for five years consecutively, from 2010-2015 [19-21]. Previous studies have reported the presence of antibodies to RVFV in different parts of Tanzania during the IEP. A study conducted by Swai and Schoolman reported the prevalence of antibodies to RVFV in Tanga region [22]. A cyclical recurrence of RVF in Tanzania has been documented whereby the disease has been recurring in approximately ten-year cycles with three major epidemics reported in 1977, 1997/98 and 2006/2007 [6]. The areas affected by the epidemics have been reported to vary whereas RVF epidemics of 1977 and 1997/98 were mainly confined to northern regions of Tanzania, the 2006/2007 outbreak assumed a larger scale involving 10 out of the 21 regions of Tanzania with human cases reported in Arusha, Dar es Salaam, Dodoma, Iringa, Manyara, Mwanza, Morogoro, Pwani, Singida and Tanga Regions [23]. Studies have not been able to detect the viruses in its hosts (mosquitoes, ruminants and human hosts) during IEPs.

Despite the failure to detect RVFV by PCR, we report a significant seroprevalence of 16.39% for RFV antibodies in the Wami-Dakawa site. This confirms the active circulation of RVFV in Wami-Dakawa area. The ecology of the Wami-Dakawa area and life sustaining economic activities (agropastoralism) of the residents of Wami-Dakawa are among the probable environments that offers suitable conditions for maintenance of RVFV in the area. Despite the positive serological results for RVFV, this study did not diagnose any RVFV among participants by PCR. This raises the unanswered question of how the viruses are maintained across their hosts during inter epidemic periods. A clear definition of the transmission would constitute a prime target for RVF and other vector borne viral infections. We detected none of the studied viruses by PCR, and observed an absence of seropositive individuals to any of the viruses in the Magugu site. Interesting as this observation may be, the relative “sterility” of the Magugu site to mosquito borne infections warrants an epidemiological explanation owing to the otherwise conducive environment for arboviral transmission including the presence of suitable climate and breeding sites for mosquitoes. Previous studies on mosquito borne infections in Magugu had reported a similar state of sterility for mosquito borne infections for the past decade [14, 24-27].

Recent epidemiological data on CHIKV and DENV outbreaks in Tanzania has largely been reported in Dar-es Salaam the capital city of Tanzania [19] and Tanga region [15]. There have been no reported DENV and CHIKV epidemics since the reported 2014-2015 outbreaks. Recently in 2019, a DENV outbreak occurred involving largely Dar es Salaam and its neighboring regions of Coast and Morogoro regions, and Tanga region Morogoro (https://www.iamat.org/country/tanzania/risk/dengue). Despite the presence of reported cases of DENV in Morogoro urban a few months prior to this study (Unpublished), our study could not detect any seropositive individuals in Wami-Dakawa. This finding was unexpected due the closeness of Wami-Dakawa and Morogoro urban that includes a rich day to day human interactions. This can be associated with mosquito abundance and human-human interaction during the dry season (August) of the year. These are influenced with farming activities, harvesting and weather conditions. Since we have not reported presence of the viruses in Aedes mosquitoes in the current study, an important avenue for a future study would be to conduct an entomological survey that would establish vector abundance and DENV infection rates.

In this study, there was a higher proportion of individuals seropositive to RVFV (16.39%) than individuals seropositive to CHIKV (9.83%), although the difference was not statistically significant. While RVFV has non-human domestic animals as one of its hosts, CHIKV and DENV do not infect livestock, making the former easier to maintain among high risk communities, such as livestock herders, during IEPs. Sampling for this study was conducted during the dry season. It is worthwhile to recommend that this variation is followed up in tandem with entomological screening, across different seasons of the year to get an insight of how infection and seroconversion may vary as a means of RVFV, DENV and CHIKV maintenance in the absence of outbreaks.

## Conclusion and Recommendation

We report antibodies to RVFV to be the most prevalent followed by antibodies to CHIKV in Wami-Dakawa, in Mvomero district of Morogoro region during dry season. This study did not detect any individual in Wami-Dakawa who was seropositive to DENV. Magugu site was found to be free from both infection and exposure to RVFV, CHIKV and DENV. Larger numbers of household members in a house, having no formal education and having recently travelled to an urban destination were risk factors being seropositive to RVFV and CHIKV. Since arbovirus outbreaks occur sporadically and usually unpredictable in nature, it is crucial to undertake active surveillance measures for RVFV, DENV, CHIKV and other viral agents in endemic countries. This strategy is urgent in the present time characterized with drastic climatic changes.

## Acknowledgement

This study was conducted financial support from the International Center for Genetic Engineering and Biotechnology (ICGEB), Grant #CRP/TZA18-04 and The East African Consortium for Clinical Research (EACCR-2) grant # EDCTP-RegNet2015-1104T through the KCRI NID node. This support is highly acknowledged. We acknowledge the logistical support provided by Executive officers in the study sites, participants who consented to be part of this study and the administrative team at KCRI.

